# An Analysis of Protein Language Model Embeddings for Fold Prediction

**DOI:** 10.1101/2022.02.07.479394

**Authors:** Amelia Villegas-Morcillo, Angel M. Gomez, Victoria Sanchez

## Abstract

The identification of the protein fold class is a challenging problem in structural biology. Recent computational methods for fold prediction leverage deep learning techniques to extract protein fold-representative embeddings mainly using evolutionary information in the form of multiple sequence alignment (MSA) as input source. In contrast, protein language models (LM) have reshaped the field thanks to their ability to learn efficient protein representations (protein-LM embeddings) from purely sequential information in a self-supervised manner. In this paper, we analyze a framework for protein fold prediction using pre-trained protein-LM embeddings as input to several fine-tuning neural network models which are supervisedly trained with fold labels. In particular, we compare the performance of six protein-LM embeddings: the LSTM-based UniRep and SeqVec, and the transformer-based ESM-1b, ESM-MSA, ProtBERT, and ProtT5; as well as three neural networks: Multi-Layer Perceptron (MLP), ResCNN-BGRU (RBG), and Light-Attention (LAT). We separately evaluated the pairwise fold recognition (PFR) and direct fold classification (DFC) tasks on well-known benchmark datasets. The results indicate that the combination of transformer-based embeddings, particularly those obtained at amino acid-level, with the RBG and LAT fine-tuning models performs remarkably well in both tasks. To further increase prediction accuracy, we propose several ensemble strategies for PFR and DFC, which provide a significant performance boost over the current state-of-the-art results. All this suggests that moving from traditional protein representations to protein-LM embeddings is a very promising approach to protein fold-related tasks.

## 1 Introduction

Despite recent breakthroughs in predicting protein three-dimensional structures with high accuracy (AlphaFold [1, 2] and RoseTTAFold [3]), there is still a special interest in identifying the fold type of a protein. This allows for a better understanding of the functionality of the newly solved structures (e.g. AlphaFold DB [4]), and it is often accomplished by classifying them according to structural and sequential similarities with respect to known proteins. In this regard, structural classification databases such as SCOP [5, 6] and CATH [7, 8] already provide a hierarchical grouping of protein domains from the protein data bank (PDB) [9, 10] into different categories. In SCOP these are named structural class, fold, superfamily and family, with increasing amino acid sequence similarity at each level. Among them, the focus is on obtaining accurate predictions at the fold level, where protein domains have a similar arrangement of structural elements but substantially differ in the amino acid sequence, a problem widely known as protein fold recognition [11–15].

During the past few decades, many computational methods have been proposed to predict the fold class of a protein domain. These can be divided according to the task they aim to solve. The first one is *pairwise fold recognition* (PFR), in which the fold class of the query protein is inferred by comparing with templates with known structure [16–18]. PFR approaches mainly include methods based on homology modeling (sequence alignments [19], profile alignments [20], and Markov random fields [21]); threading [22–28]; machine learning for binary classification [29–31]; multi-view learning [32–34]; and learning to rank [35–37]. Another set of methods use deep learning to learn *fold-representative embeddings*, which are then used to measure the structural similarity between two protein domains. DeepFR [38] introduced this methodology, which has been followed by more recent approaches [39–45] using either predicted contact maps, or evolutionary information as input representation of the protein. The second task is *direct fold classification* (DFC), in which the protein sequences are directly mapped into a pre-defined group of fold classes [46]. Most of the proposed methods [47–56] had used evolutionary information and machine learning to classify only a small portion of all possible SCOP folds (i.e. the most populated ones). In contrast to them, DeepSF [57] was the first method to perform fold classification into one of the more than thousand existing folds in SCOP through deep learning.

Traditionally, the state-of-the-art methods for different protein-related tasks have used evolutionary information in the form of multiple sequence alignment (MSA) as input source. Following a more modern approach, recent methods use protein representations that are extracted from pre-trained protein language models (protein-LMs). These models have been taken from the field of NLP (natural language processing) [58], by treating the protein sequences as sentences, and the amino acids as word equivalents. More specifically, the models learn meaningful representations of the proteins (*protein-LM embeddings*) in a self-supervised manner [59] by using the vast amount of unlabeled sequences contained in protein databases such as Swiss-Prot [60], Pfam [61], and UniRef [62] (all based on UniProt [63]); and metagenomic databases such as the big fantastic database (BFD) [64, 65]. This way, ProtVec [66], based on word2vec [67, 68], was the first method proposed to extract protein representations from their sequences. Unlike word2vec, more sophisticated protein-LMs take into account both the context and order of amino acids in the sequence through LSTM (long short-term memory) recurrent units [69]. These include methods such as UDSMProt [70] and UniRep [71], using weight-dropped LSTMs and multiplicative LSTMs, respectively; as well as two methods based on the ELMo model [72]: SeqVec [73] and the language model part from [74]. The next generation of protein-LMs come from the use of transformer architectures based on self-attention mechanisms [75]. Examples are TAPE-Transformer [76], ESM-1b [77], ESM-MSA [78] and ProtBERT [79], which have been inspired in the BERT model [80]. In addition to BERT, the ProtTrans project [79] explored the use of other five transformer architectures [81–85] for protein representation learning. These pre-trained models allow for transfer learning to different protein-related downstream tasks in which the amount of labeled sequences in the databases is significantly smaller. By using simple supervised models, protein-LM embeddings have proven to be successful in predicting protein secondary structure, subcelullar localization, and remote homologs [73, 76, 79, 86], as well as protein function [87–89] and sequence variation [90, 91]. It has been also found that the attention layers in transformer models are able to learn protein contact map information directly from self-supervised training on sequences [77, 78, 92].

Our proposal here (summarized in Figure 1) is to leverage several pre-trained protein-LM embeddings for fold prediction. We hypothesize that self-supervised training of the protein-LMs might capture fold information from millions of protein sequences, and therefore the learned representations could be useful for comparison of structurally similar proteins (PFR task), as well as classification into fold classes as defined by SCOP (DFC task). To test this, we followed the same idea of the DeepSF, DeepFR, and subsequent deep learning models for fold prediction. These neural network models allow for the fine-tuning of protein-LM embeddings (here *LMEmb*) to learn new fold-representative embedding vectors (here *FoldEmb*), while performing fold classification. In both tasks, PFR and DFC, we compared the performance of different neural network architectures working on either amino acid-level or protein-level embeddings. Comparison with previous methods also allowed us to analyze the impact of changing traditional protein evolutionary features by protein-LM embeddings as input representations. All in all, we found that transformer-based protein-LM embeddings are particularly useful for protein fold prediction, outperforming the state-of-the-art results for both fold recognition and fold classification.

**Figure 1:**
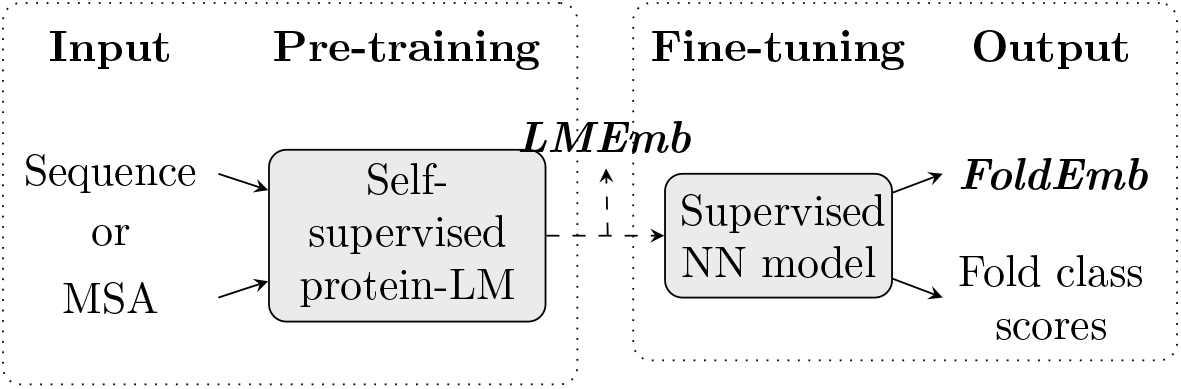
Overview of our approach for protein fold prediction. First, we extract a protein embedding representation from the amino acid sequence or multiple sequence alignment (MSA) using protein language models (protein-LMs) which have been pre-trained in a self-supervised manner (i.e. using the input sequential information itself). As a result, we obtain a protein-LM embedding (*LMEmb*) of size *L* × *F*, where *L* is the length of the protein sequence and *F* is the size of the amino acid-level embedding. Then, we fine-tune this embedding through a neural network (NN) model that is trained, in a supervised manner, to map the input protein into *K* fold classes. The outputs of this model are, on the one hand, a fold-representative embedding of the protein (*FoldEmb* with fixed-size 512), used to perform the pairwise fold recognition (PFR) task; and, on the other hand, the scores for each fold class, used in the direct fold classification (DFC) task.

## 2 Materials and Methods

### 2.1 Input Protein Information

Our framework for fold prediction (Figure 1) takes as input sequential information of the protein, either the amino acid sequence or a multiple sequence alignment (MSA). To build the MSAs for our protein domains, we followed the pipeline specified in [78]. That is, we first generated the MSA by running HHblits [93] against the uniclust30_2017_10 database [94], with number of iterations equal to 3. The resulting MSA was then subsampled by filtering the number of sequences down to 256 with hhfilter [93]. If more than 256 sequences were returned, we applied the diversity maximizing strategy from [78] to select those sequences with highest average hamming distance.

### 2.2 Pre-Trained Protein-LM Embeddings

As protein representations, we used self-supervised embeddings from pre-trained protein language models (protein-LMs). We analyzed the performance of LSTM-based protein-LMs such as UniRep [71] and SeqVec [73], as well as several transformer-based models such as ESM-1b [77], ESM-MSA-1b (here ESM-MSA) [78], ProtBERT-BFD and ProtT5-XL-U50 (here ProtBERT and ProtT5) [79]. We denote these self-supervised embeddings as *LMEmb*, in order to differentiate them from our fine-tuned embeddings, *FoldEmb*, which are supervisedly trained using fold labels. The total size of an *LMEmb* embedding for a protein of length *L* is *L* × *F*, where *F* is the size of the individual embedding provided by the protein-LM for each amino acid. By averaging the embedding matrix over the length dimension, we can obtain an alternative fixed-size representation for the protein domain (size *F*). We will refer to this representation as protein-level embeddings or *LMEmb-Prot* (size *F*) in contrast to the amino acid-level embeddings defined above and denoted as *LMEmb-AA* (size *L* × *F*). Table 1 summarizes the training details for each protein-LM embedding included in the analysis.

**Table 1:** Characteristics of the protein language model (protein-LM) embeddings used in this analysis.

| Embeddings | Size ($F$) | Language Model | #Layers | #Parameters | Training Database |
| --- | --- | --- | --- | --- | --- |
| UniRep [71] | 1900 | mLSTM | 1 | 18M | UniRef50 (24M seqs) |
| SeqVec [73] | 1024 | ELMo (BLSTM) | 2 | 93M | UniRef50 (33M seqs) |
| ESM-1b [77] | 1280 | BERT (Transformer) | 33 | 650M | UniRef50 (27M seqs) |
| ESM-MSA [78] | 768 | BERT (Transformer) | 12 | 100M | UniRef50 (26M MSAs) |
| ProtBERT [79] | 1024 | BERT (Transformer) | 30 | 420M | BFD (2B seqs) |
| ProtT5 [79] | 1024 | T5 (Transformer) | 24 | 3B | BFD (2B seqs) + UniRef50 (45M seqs) |

#### LSTM-Based Models

UniRep [71] and SeqVec [73] are two protein-LMs trained using recurrent layers with long short-term memory (LSTM) [69] units. Both models were trained in an auto-regressive manner, trying to predict the next amino acid given all previous amino acids in a protein sequence. The *UniRep* model consists of one layer of multiplicative LSTM (mLSTM) [95] with 1900 hidden units, trained on 24 million protein sequences from the UniRef50 database. We used the TAPE [76] implementation^1^ and the pre-trained model from UniRep to extract *L ×* 1900 dimensional embeddings for our protein domains. On the other hand, *SeqVec* is based on the ELMo model [72] from NLP. The SeqVec model was trained on 33 million protein sequences from UniRef50. Its architecture is formed by one CharCNN layer to embed the input characters (amino acids), followed by two layers of bidirectional LSTMs (BLSTM) [96] with shared parameters for the forward and backward passes. We obtained *L ×* 1024 dimensional SeqVec embeddings by concatenating both directions of the LSTMs and then adding the outputs of the three layers. To do so, we used the official code^2^ and the pre-trained model of SeqVec.

#### Transformer-Based Models

We consider two sets of transformer-based protein-LMs—evolutionary scale modeling (ESM) [77, 78] and ProtTrans [79]. The ESM models are based on the BERT transformer architecture [80]. Unlike auto-regressive LSTM-based protein-LMs, ESM models were trained to predict masked amino acids using all preceding and following amino acids in the sequence. This training objective is referred to as masked language modeling (MLM). In our analysis, we consider the pre-trained *ESM-1b* model [77], which has 33 transformer layers and a total of 650M parameters, and was trained on 27 million protein sequences from UniRef50. Instead of individual protein sequences, the *ESM-MSA* model [78] was trained on multiple sequence alignments (MSAs) constructed from sequences in UniRef50 (26 million of MSAs). This model uses the axial attention mechanism from [97] and has fewer transformer layers (12) and parameters (100M) than ESM-1b.

To obtain an embedding for each protein domain in our datasets we used the official code^3^ and the pretrained ESM-1b and ESM-MSA models. For ESM-1b, we extracted amino acid-level embeddings from the last transformer layer, resulting in *L ×* 1280 vectors. In contrast, the ESM-MSA model provides embeddings for each sequence in the MSA, so the output of the last transformer layer is of size 256 *× L ×* 768. We averaged over all sequences in the MSA to obtain final embeddings of size *L ×* 768.

In contrast to the ESM models, the ProtTrans project [79] scaled up the transformer-based protein-LMs to leverage metagenomic databases with billions of protein sequences, leading to architectures with several billions of parameters. In this work, we consider models based on two auto-encoder transformers, BERT and T5 [83], denoted here as ProtBERT and ProtT5 respectively. The *ProtBERT* model has 30 layers and a total of 420M parameters, and has been trained on 2 billion protein sequences from the BFD database. Unlike ProtBERT and the previous ESM models, which only train the encoder component, *ProtT5* includes both the encoder and decoder, with 24 layers and a total of 3B parameters. It was trained on the BFD database and later refined using 45 million sequences from UniRef50. To extract *L ×* 1024 dimensional embeddings for our protein domains, we used the official ProtTrans implementation^4^ and the pre-trained models for ProtBERT and ProtT5. Following the recommendations in [79], we only used the encoder part of ProtT5 to embed our protein domains.

### 2.3 Neural Network Models for Protein Embedding Fine-Tuning

In this subsection we describe the neural architectures and the corresponding supervised training procedures for fine-tuning the protein-LM embeddings *LMEmb* into the fold representative embeddings *FoldEmb* (see Figure 1).

#### 2.3.1 Neural Architectures

Our neural network models include a supervised embedding extractor followed by a linear classifier. Specifically, the embedding extractors accept as input either the amino acid-level protein-LM embeddings *LMEmb-AA* or the averaged protein-level embeddings *LMEmb-Prot*. We used three different neural architectures to extract supervised embeddings, *FoldEmb*, of fixed-size (512): a Multi-Layer Perceptron (MLP), our previously proposed Residual-Convolutional network and Bidirectional Gated Recurrent Unit (RBG) [45], and the Light-Attention (LAT) architecture from [86]. These architectures are illustrated in Figure 2. For the last step classifier we use a single fully-connected (FC) layer which projects the *FoldEmb* embeddings into *K* output elements (i.e. logits) corresponding to the fold classes. The supervised embedding extractor and classifier are trained together in an end-to-end fashion.

**Figure 2:**
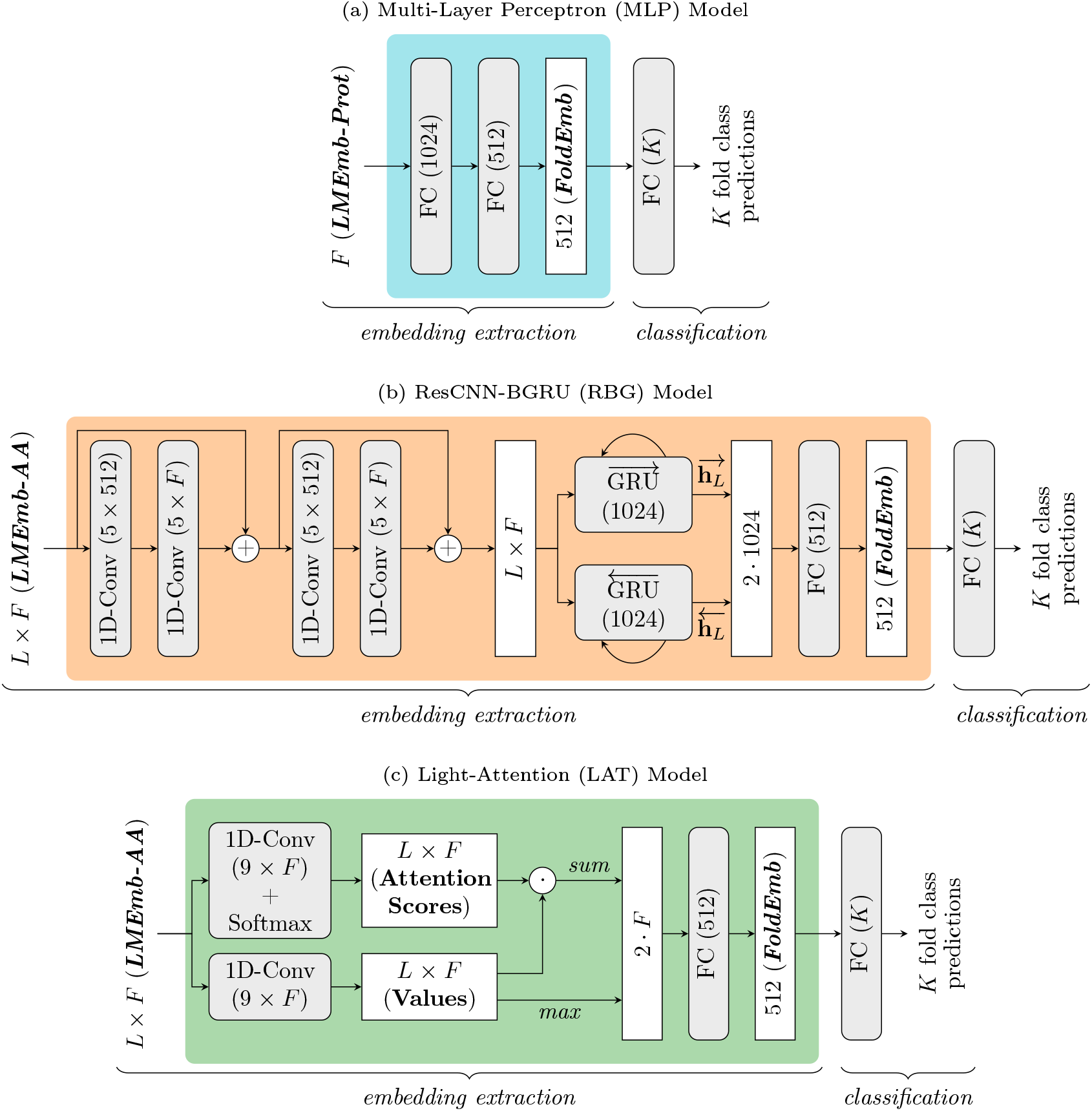
Neural network models used to fine-tune the protein-LM embeddings (*LMEmb*) to fold-representative embeddings (*FoldEmb*), as well as to perform direct fold classification. Three neural architectures are used as embedding extractors (identified with three distinct background colors). **(a)** The Multi-Layer Perceptron (MLP) model processes the protein-level embeddings (*LMEmb-Prot*) through two fully-connected (FC) layers. **(b)** The ResCNN-BGRU (RBG) model [45] processes the amino acid-level embeddings (*LMEmb-AA*) through two residual-convolutional blocks, a bidirectional gated recurrent unit (GRU) layer, and an FC layer. **(c)** The Light-Attention (LAT) model, adapted from [86], also processes *LMEmb-AA* through an attention mechanism followed by an FC layer.

##### Multi-Layer Perceptron (MLP)

As a reference model, we processed the *LMEmb-Prot* embeddings through a simple MLP. The architecture consists of two FC layers with 1024 and 512 neurons each (see Figure 2a) with ReLU activation. After each layer, we also apply batch-normalization [98] and dropout [99] with drop probability *p* = 0.5 to prevent overfitting during training.

##### Residual-Convolutional Network and Bidirectional Gated Recurrent Unit (RBG)

To process *LMEmb-AA* embeddings, we consider our ResCNN-BGRU (RBG) model architecture (Figure 2b) which obtained state-of-the-art performance on protein fold recognition [45]. This architecture consists of three distinct parts, which we briefly describe here (further details can be found in [45]). First, the residual-convolutional part consists of two identical residual blocks with skip connections. Each block contains two 1D-convolutions with 512 and *F* filters of length 5, where *F* is the dimensionality of the input embedding (see Table 1). The 1D-convolutions are followed by ReLU activation, batch-normalization and dropout (*p* = 0.2). Second, a bidirectional recurrent layer is applied on top of the *L × F* outputs of the residual-convolutional part. This consists of a bi-directional gated recurrent unit (GRU) [100] layer with 1024 state units for each direction. To obtain a fixed-size vector, we concatenate the last states from both forward 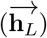 and backward 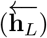 GRU layers into a vector of 2048 elements. Finally, we project this vector into a 512-dimensional embedding (*FoldEmb*) using an FC layer of size 512, followed by hyperbolic tangent (tanh) activation and dropout (*p* = 0.2).

##### Light-Attention (LAT)

We also include in our analysis the Light-Attention (LAT) model from [86], which has been recently proposed for predicting protein subcellular location, using *LMEmb-AA* embeddings as inputs. The architecture is shown in Figure 2c. It applies two parallel 1D-convolutions with *F* filters of length 9, to produce the attention coefficients and value features separately. The attention weights are obtained from the coefficients by applying the softmax operation over the length dimension. The resulting weights are used to compute a weighted sum of the values, producing an *F* -dimensional vector independent of the protein length. This vector is then concatenated with the max-pooled values (across the length dimension) to produce a vector of size 2*F*. To compute the 512-dimensional *FoldEmb* embeddings, we adapted the MLP part of the original LAT architecture by including an FC layer similar to that described above for the RBG model.

#### 2.3.2 Model Optimization

The fine-tuning models were trained to minimize the *large margin cosine loss* (LMCL) [101] between the predicted and true fold classes for each protein domain in the training dataset. The LMCL is an *L*_2_-normalized and margin discriminative version of the softmax cross-entropy loss. The *L*_2_ normalization enforces the *FoldEmb* vectors to be distributed on the surface of a hypersphere. It is formally defined as:

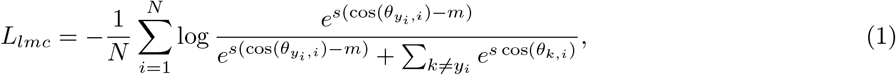

where *N* is the number of training samples in the mini-batch and *K* is the number of fold classes. The cosine is computed as 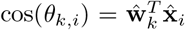 where ŵ_*k*_ and 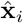 are *L*_2_-normalized versions of the *k*-th class weight vector **w**_*k*_ (from the last classification layer, bias is set to zero for simplicity) and the *i*-th embedding vector **x**_*i*_ (here the 512-dimensional *FoldEmb*). As can be noticed from Eq. 1, this loss introduces two hyperparameters, the scale and margin (*s, m ≥* 0). The scaling hyperparameter *s* controls the radius of the hypersphere on which the embeddings are distributed, while the margin *m* controls the decision boundaries between fold classes, and so the capacity of learning more discriminative embeddings. Following previous results [45] and light hyperparameter tuning using cross-validation, we use the same scale for all our networks (*s* = 30) and a margin with value *m* = 0.2 for the MLP model and *m* = 0.6 for the models working on *LMEmb-AA* embeddings (that is, RBG and LAT).

For training, we use mini-batches with 64 protein domains each, and the Adam optimizer [102] with an initial learning rate equal to 10^−3^. To prevent overfitting, along with the batch-normalization and dropout techniques specified before, we apply *L*_2_ penalty with a weight decay of 5 · 10^−4^. All models were trained for 80 epochs and we decreased the learning rate by a factor of 10 at epoch 40, as it proved to improve the performance in previous works [44, 45]. We implemented our models using PyTorch [103] and executed them on a single GPU (NVIDIA Tesla V100) with 32GB of memory.

### 2.4 Evaluation Tasks

This subsection details the scoring procedures, ensembling strategies, and performance metrics used to evaluate the two protein fold-related tasks: pairwise fold recognition (PFR) and direct fold classification (DFC). As a reference, an evaluation where the protein-LM embeddings are directly used without fine-tuning (i.e. without additional supervised training) was also considered for both tasks.

#### 2.4.1 Pairwise Fold Recognition (PFR)

##### PFR Task

The pairwise fold recognition task involves evaluating the structural similarity of two protein domains. To do so, we used the 512-dimensional supervised embeddings (*FoldEmb*) extracted from our neural network models (Figure 2). These embeddings were used to compute a fold similarity score for each of two domains within the test set, indicating whether they belong to the same fold class or not. Following previous works, we used the cosine similarity as the similarity metric [38, 44, 45].

##### Ensemble Strategies

Since ensemble methods have been found to be promising for PFR in previous works [38, 43–45], we also leveraged ensembling techniques here to obtain a better fold similarity score from our best performing *FoldEmb* embeddings. The first type of ensembling strategy, which we refer to as *average ensemble*, involves directly averaging the cosine similarity scores, provided by the chosen models, for each pair of protein domains in the test set. The second ensembling strategy consists of training a random forest (RF) model using our cosine similarity scores along with the 84 pairwise similarity measures from [30, 31]. We refer to this strategy as *random forest ensemble*. The RF model was trained to determine whether the protein domains in each pair share the same fold class using the vector of pairwise scores as input, and a total of 500 decision trees. Note that this strategy involves training an RF model, so we evaluated using a 10-stage cross-testing setting over the test set as in [38, 44, 45].

##### Ranking and Evaluation

To evaluate an individual protein domain (query), we ranked the rest of domains in the test set (templates) by fold similarity score, and then assigned the fold class of the most similar one to the query. As originally proposed in [13] and following subsequent works [38, 44, 45], this evaluation was performed at three levels with increasing difficulty, according to the SCOP hierarchy—*family, superfamily* and *fold* [5]. At each level, a positive pair contains two protein domains sharing the same class in the selected level, but a different class in the level immediately below. For example, at the superfamily level, two domains within a positive pair belong to the same superfamily class (and therefore the same fold class), but different family classes. Irrespective of the level, all negative pairs were evaluated, each one containing two protein domains from different fold classes. After the ranking and fold assignment, we computed the ratio of hits (top 1 accuracy), as well as the ratio of finding the correct fold class within the 5 top-ranked templates (*top 5 accuracy*).

#### 2.4.2 Direct Fold Classification (DFC)

##### DFC Task

In the direct fold classification task, we evaluated the ability of our neural network-based models to classify the input test domains into *K* fold classes. This task was originally proposed in [57]. In this case, instead of extracting the supervised embeddings (*FoldEmb*), we obtained a score (i.e. logit) for each fold class from the last classification layer of our models (Figure 2), and the predicted fold as the one maximizing the class scoring vector.

##### Ensemble Strategies

As with the PFR task, we combined the best performing fine-tuning models to generate several ensembles. In this case, we followed the *soft voting ensemble* approach to get a better prediction for each tested protein domain. This strategy involves computing the vector of logits from each model and accumulating them in a new class scoring vector. Then, we assign the fold which maximizes this scoring vector for each query domain.

##### Evaluation

We also assessed the DFC task at the *family, superfamily* and *fold* levels. In contrast to the PFR task, now the full test set is split into three subsets, each one containing test domains that only share the specific level with domains from the training dataset. As performance metric, we consider the ratio of protein domains that were correctly classified (*top 1 accuracy*), as well as the ratio of finding the correct fold within the 5 top-scoring classes (*top 5 accuracy*).

#### 2.4.3 Baseline Evaluation of Protein-LM Embeddings

To provide a baseline comparison, we also evaluated the PFR and DFC tasks directly with the pre-trained protein-LM embeddings (*LMEmb*). We used the cosine similarity metric for the protein-level embeddings *LMEmb-Prot*. In contrast, for the amino acid-level embeddings *LMEmb-AA*, we computed the soft symmetric alignment (SSA) proposed in [74] but using cosine similarity instead of *L*_1_ distance. That is, given two sequences of embeddings 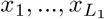 and 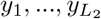, SSA is obtained as:

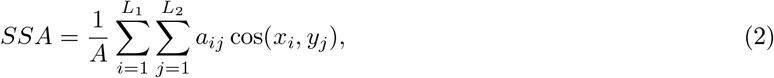

where 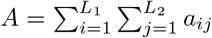 acts as a normalization factor, and *a*_*ij*_ = *α*_*ij*_ + *β*_*ij*_ *− α*_*ij*_*β*_*ij*_ represent the alignment matrix, whose components are computed as:

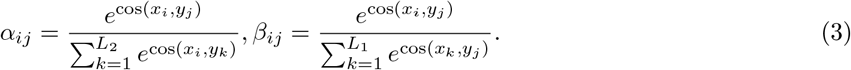

Note that, while the baseline PFR task is performed using pairs from the test set only, the DFC one involves computing the fold similarity metric for each test embedding against all the training ones, and then assigning the fold class of the closest training domain.

### 2.5 Datasets

To assess the aforementioned tasks, we trained the fine-tuning models using 16133 protein domains from SCOPe version v2.06 [38], which are classified into *K* = 1154 folds. For PFR, we tested the models using the well-known LINDAHL dataset [13] containing 976 protein domains from 330 distinct folds. For the DFC task, we used the updated version of LINDAHL to SCOP v1.75 [44] (named LINDAHL_1.75), where, in order to directly classify the test domains, we keep only those (of the 976) that share their fold class with one of the *K* seen during training. This resulted in a test set with 871 domains from 244 folds.

In addition, for DFC we also evaluated the performance over the SCOP_2.06 test set [57], which contains 2533 protein domains from 550 folds. To avoid overlap between this test set and the training set described above (both are derived from SCOPe v2.06), in this particular case we used the training set proposed in [57], which contains 16712 domains classified into *K* = 1195 folds (from SCOP v1.75).

For each task and test set, Table 2 summarizes the number of individual protein domains and distinct folds evaluated at each level (family, superfamily and fold). In all cases, the protein domains within each training set share at most 95% sequence similarity. Moreover, the maximum sequence identity within each of the test sets is 40%, as well as with respect to their respective training sets.

**Table 2:** Number of protein domains and fold classes evaluated in each test set.

| Task | Test Set | Full Set |  | Family Level |  | Superfamily Level |  | Fold Level |  |
| --- | --- | --- | --- | --- | --- | --- | --- | --- | --- |
|  |  | #Domains | #Folds | #Domains | #Folds | #Domains | #Folds | #Domains | #Folds |
| PFR | LINDAHL | 976 | 330 | 555 | 121 | 434 | 79 | 321 | 38 |
| DFC | LINDAHL_1.75 | 871 | 244 | 591 | 177 | 210 | 107 | 70 | 37 |
|  | SCOP_2.06 | 2533 | 550 | 742 | 316 | 1754 | 418 | 37 | 15 |

The LINDAHL test set is evaluated on the pairwise fold recognition (PFR) task, whereas the LINDAHL\_1.75 and SCOP\_2.06 test sets are evaluated on the direct fold classification (DFC) task. We also provide the number of domains and folds evaluated at the family, superfamily and fold levels.

## 3 Results and Discussion

### 3.1 Performance of Self-Supervised *LMEmb* Embeddings in PFR and DFC Tasks

We first evaluated how the self-supervised *LMEmb* embeddings perform in predicting structural similarity using the LINDAHL test set. In Table 3 we provide the pairwise fold recognition (PFR) results for the two types of embeddings, *LMEmb-AA* and *LMEmb-Prot*, from the 6 protein-LMs we considered. We find that aggregating embeddings across the protein length (*LMEmb-Prot*) helps in all cases when using cosine similarity. Note that, as can be seen in Table S1 (Supplementary Material), using *L*_1_ distance as comparison metric shows similar results to cosine similarity for all *LMEmb-Prot* embeddings.

**Table 3:** Performance of the *LMEmb* embeddings in the pairwise fold recognition (PFR) task, using the LINDAHL test set.

| Embeddings | Family |  | Superfamily |  | Fold |  |
| --- | --- | --- | --- | --- | --- | --- |
|  | Top 1 | Top 5 | Top 1 | Top 5 | Top 1 | Top 5 |
| ***LMEmb-AA* ($L \times F$)** |  |  |  |  |  |  |
| UniRep | 28.5 | 49.2 | 25.3 | 38.9 | 14.3 | 35.8 |
| SeqVec | 48.3 | 66.5 | 27.2 | 47.0 | 13.7 | 29.3 |
| ESM-1b | 7.4 | 14.8 | 2.1 | 6.7 | 0.6 | 8.1 |
| ESM-MSA | 27.9 | 47.7 | 13.8 | 29.5 | 12.1 | 22.4 |
| ProtBERT | 18.9 | 34.8 | 4.4 | 15.9 | 4.4 | 15.9 |
| ProtT5 | 76.4 | 87.4 | 34.3 | 56.5 | 14.0 | 29.9 |
| ***LMEmb-Prot* ($F$)** |  |  |  |  |  |  |
| UniRep | 45.6 | 60.9 | 35.0 | 47.7 | 19.3 | 35.8 |
| SeqVec | 62.3 | 77.8 | <u>44.9</u> | 60.6 | 18.4 | <u>37.7</u> |
| ESM-1b | <u>81.6</u> | 88.5 | <u>44.9</u> | 59.7 | <u>21.2</u> | 37.1 |
| ESM-MSA | 76.6 | 88.1 | 42.9 | 54.4 | 16.2 | 28.0 |
| ProtBERT | 42.9 | 58.6 | 10.8 | 27.2 | 8.4 | 22.4 |
| ProtT5 | 81.1 | <u>90.8</u> | 40.3 | <u>62.0</u> | 16.5 | 32.4 |

The top 1 and top 5 accuracy (%) results are provided at the family, superfamily and fold levels. We compare the performance of the amino acid-level embeddings *LMEmb-AA* (using the SSA metric in Eq. 2) and protein-level embeddings *LMEmb-Prot* (using the cosine similarity metric). Underline indicates best performance.

Amongst the *LMEmb-Prot* embeddings, the ESM-1b ones yield better overall PFR results, while the Prot-BERT embeddings perform the worst at the three levels. We also notice differences across levels. For example, the ESM-1b and ProtT5 embeddings perform the best at the family level, with a high accuracy (81% top 1). This suggests that these embeddings could be used for homology searching when the amino acid sequence similarities are high. Furthermore, ESM-1b outperforms the rest of embeddings at the fold level (21.2%). Nevertheless, the accuracy values are generally low at this level. A similar trend is observed in Table S2 (Supplementary Material) for the direct fold classification (DFC) task evaluated on the LINDAHL_1.75 and SCOP_2.06 test sets. It should be noted that the overall poor performance of protein-LM embeddings at the fold level is to be expected, as they have been learned in a self-supervised manner without any information about the fold type.

### 3.2 Performance of Fine-Tuned *FoldEmb* Embeddings in PFR Task

Figure 3 summarizes the PFR results of the fold-representative embeddings, *FoldEmb*, resulting from fine-tuning the *LMEmb* embeddings through neural network models: MLP, RBG and LAT. As a baseline, we also include the results obtained directly using the *LMEmb-Prot* embeddings (from Table 3). As can be observed, the *LMEmb* embeddings can be significantly enhanced by fine-tuning even when a simple MLP model is used for the task (*LMEmb-Prot* vs MLP in Figure 3), especially at the superfamily and fold levels. In general, after fine-tuning, the transformer-based protein-LM embeddings (ESM-1b, ESM-MSA, ProtBERT, ProtT5) show better PFR performance than the LSTM-based ones (UniRep, SeqVec). Regarding the supervised models, the RBG and LAT, both working on *LMEmb-AA*, provide better PFR results than MLP at the superfamily and fold levels. This contrasts with the results in Table 3 for *LMEmb-AA*, suggesting that the RBG and LAT models successfully exploit the protein sequence information possibly contained in these self-supervised embeddings. Thus, the top-performing model at the fold level is ProtT5 + LAT, which obtains 82.6% top 1 and 88.5% top 5 accuracy values. In contrast, our ESM-MSA + RBG model provides the best PFR results at the family and superfamily levels, with 83.2% and 81.3% top 1 accuracy, respectively. It is therefore clear that the fine-tuned *FoldEmb* embeddings are necessary to identify structural similarity in the hardest cases—when the sequence similarities are low.

**Figure 3:**
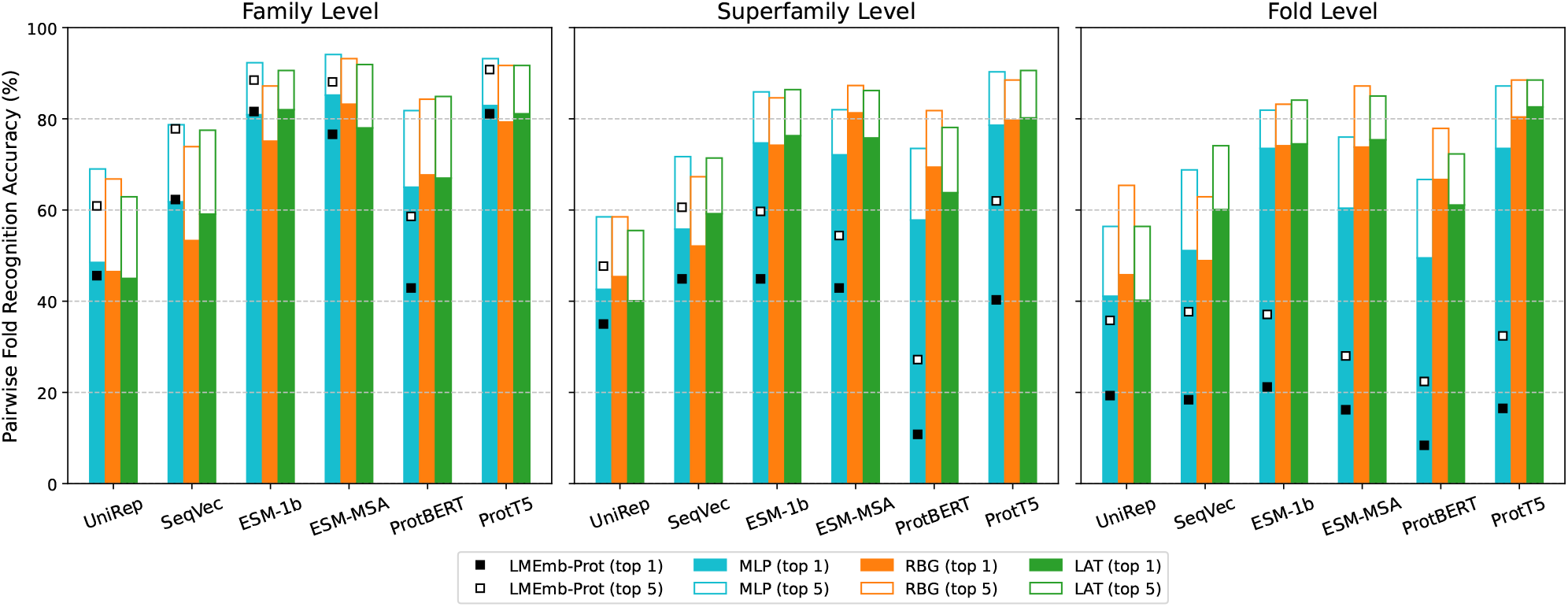
Pairwise fold recognition (PFR) accuracy (%) results on the LINDAHL test set. At each level (family, superfamily and fold), we compare the performance of the 6 protein-LM embeddings in Table 1 (UniRep, SeqVec, ESM-1b, ESM-MSA, ProtBERT, and ProtT5) fine-tuned by the 3 neural architectures from Figure 2 (MLP in cyan, RBG in orange, and LAT in green colored bars). For each one, top 1 accuracy is shown in filled bars, while top 5 accuracy is shown in empty bars. Baseline PFR results using the protein-level embeddings *LMEmb-Prot* are also included as square markers over the MLP bars (note that MLP uses these embeddings as input).

We additionally performed an ablation study (see Table S3 in Supplementary Material) using the softmax cross-entropy as loss function instead of the LMCL. This modification leads to a performance drop for most combinations of input *LMEmb* and neural architecture, which is particularly noticeable at the fold level. This confirms that the LMCL is a more suitable loss function to learn a proper organization of the fold-representative *FoldEmb* vectors in the embedding space.

### 3.3 Performance of Fine-Tuning Models in DFC Task

We evaluated the ability of our neural network models to directly classify the input protein domains into fold classes (DFC task). The classification results for the LINDAHL_1.75 and SCOP_2.06 test sets are shown in Figure 4. As in the PFR task, we also include the results provided by the *LMEmb-Prot* embeddings as a baseline (from Table S2, Supplementary Material). The results for LINDAHL_1.75 in Figure 4a show a similar pattern to those in Figure 3 for the PFR task. However, in this case, the differences in performance between the family and fold levels are more pronounced. This behavior is expected. As test domains in the family subset share a higher sequence similarity with some domains from the training set, they are likely to be easier to predict for the models. This reinforces the idea that protein-LM embeddings capture sequence similarity in proteins. By contrast, the accuracy results at the fold level are indicative of the generalization capability of the models. Here we observe that the best performing model at both the superfamily and fold levels is ProtT5 + RBG (79.5% and 55.7% top 1 accuracy, respectively), followed by ProtT5 + LAT (77.6% and 50.0%). It is also worth noting the huge differences between top 1 and top 5 accuracies of some protein-LM embeddings in the fold subset. For example, for the ESM-MSA + RBG model the top 1 accuracy is 40.0%, whereas the top 5 accuracy reaches 65.7%.

**Figure 4:**
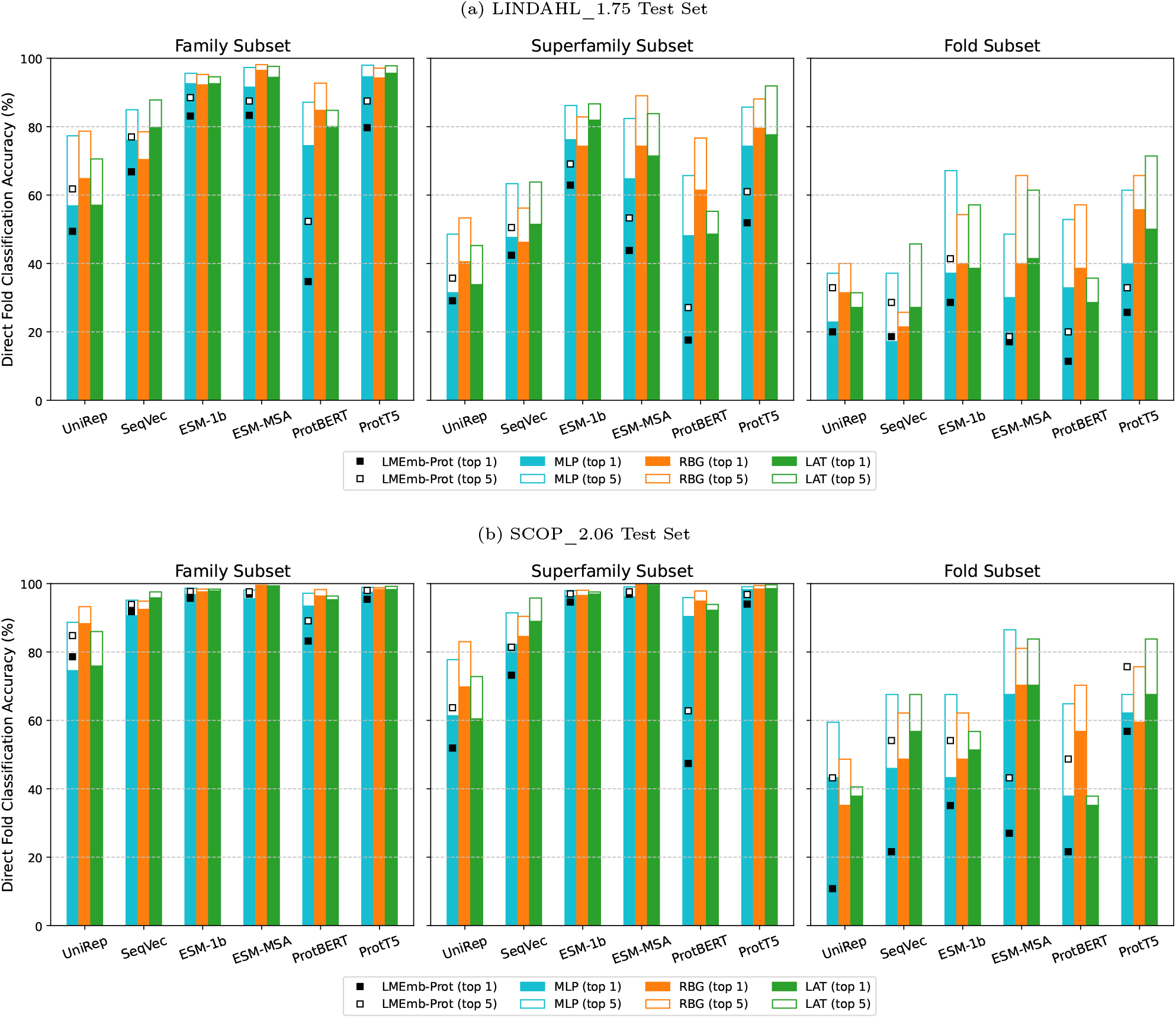
Direct fold classification (DFC) accuracy (%) results on the **(a)** LINDAHL_1.75 and **(b)** SCOP_2.06 test sets. For each subset (family, superfamily and fold), we compare the performance of the 6 protein-LM embeddings in Table 1 (UniRep, SeqVec, ESM-1b, ESM-MSA, ProtBERT, and ProtT5) fine-tuned by the 3 neural architectures from Figure 2 (MLP in cyan, RBG in orange, and LAT in green colored bars). For each one, top 1 accuracy is shown in filled bars, while top 5 accuracy is shown in empty bars. Baseline DFC results using the protein-level embeddings *LMEmb-Prot* are also included as square markers over the MLP bars (note that MLP uses these embeddings as input).

Figure 4b shows the results of the DFC task on the SCOP_2.06 test set. As can be observed, the models predict the family and superfamily levels subsets with high accuracy (close to 100% in some cases). However, similarly to the LINDAHL_1.75 test set, the fold subset in SCOP_2.06 is also difficult to classify correctly. Nevertheless, our fine-tuned models outperform the original *LMEmb* embeddings. Compared to the rest of embeddings, ESM-MSA works particularly well here for all considered neural architectures (MLP, RBG and LAT), with top 1 accuracy values around 70% at the fold level.

### 3.4 Ensemble Approaches for Increased Accuracy in PFR and DFC Tasks

Given the good performance of the transformer-based ESM-1b, ESM-MSA, and ProtT5 embeddings processed by the RBG and LAT models, we now explore the benefits of integrating them through ensemble strategies for the two assessed tasks; using average ensemble for PFR, and soft voting ensemble for DFC. Provided that the predictions from individual models are sufficiently uncorrelated, we expect an increase in performance after ensembling. In Figure 5 we show the accuracy results at the fold level for the ensembling integration of all 6 aforementioned models (the 3 protein-LMs by the 2 neural architectures). This approach outperforms other integration options that can be found in the Supplementary Material (Table S4 for PFR and Table S5 for DFC).

**Figure 5:**
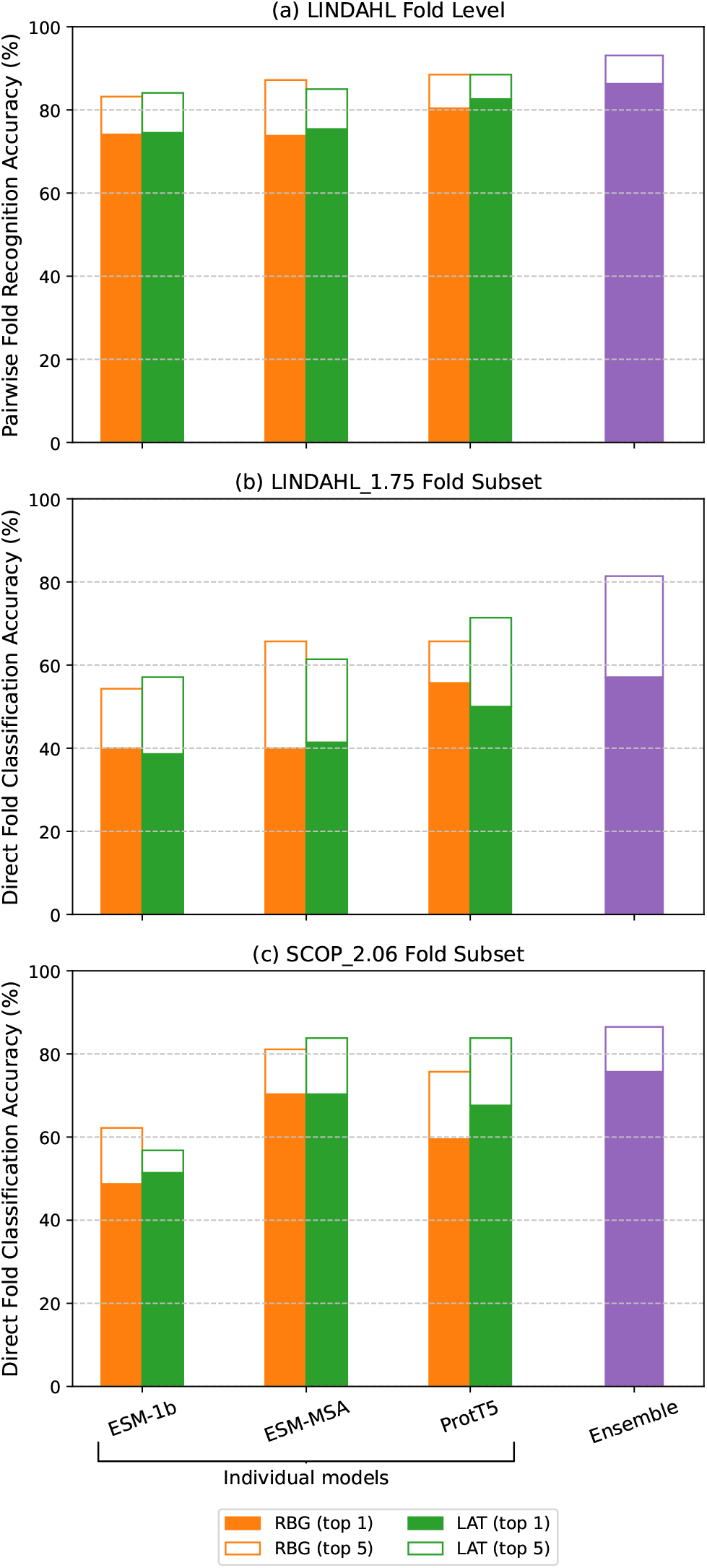
Ensemble strategy accuracy (%) results at the fold level for the **(a)** pairwise fold recognition (PFR) task using the LINDAHL test set, and the direct fold classification (DFC) task using both **(b)** LINDAHL_1.75 and **(c)** SCOP_2.06 test sets. Here the best performing ensemble strategy is shown (in purple), which involves the integration of 6 individual models. As a reference, the results of such models are also included—the 3 protein-LM embeddings (ESM-1b, ESM-MSA, and ProtT5) with neural architectures RBG (in orange) and LAT (in green). Top 1 and top 5 accuracies for each technique are represented in filled and empty bars, respectively.

As a reference, we also include the performance of the 6 individual models used in the ensemble.

For the PFR task, the LINDAHL test set is used to evaluate the ensemble strategy (Figure 5a). By simply averaging the cosine similarity scores we obtain 86.3% top 1 accuracy (93.1% top 5) at the fold level, almost 4 percentage points over the best individual model (ProtT5 + LAT). For the DFC task evaluated on LINDAHL_1.75 (Figure 5b), the ensemble approach obtains 57.1% top 1 accuracy at the fold level, slightly better than the best individual ProtT5 + RBG model (55.7%). This suggests that accurate classification at the fold level is still difficult even after ensembling. However, a noticeable improvement is observed when considering the top 5 accuracy (81.4% over the 65.7% in ProtT5 + RBG). Additionally, the ensemble approach yields better performance for the SCOP_2.06 test set (Figure 5c) in terms of both top 1 and top 5 accuracy (75.7% and 86.5%, respectively), that is, more than 5 percentage points over the best individual model (ESM-MSA + RBG).

When taking into account the family and superfamily levels (Tables S4 and S5, Supplementary Material), a performance boost over the individual models is also achieved by the ensembling approach in both tasks (PFR and DFC).

### 3.5 Comparison with State-Of-The-Art Methods for Fold Recognition and Fold Classification

Finally, we compare the performance of our best individual models, as well as the ensemble strategy we propose, against the state-of-the-art results for fold recognition and fold classification. First, we compare to several methods intended for the PFR task, which can be grouped into three categories: (i) alignment and threading methods such as PSI-BLAST [104], HHpred [20], RAPTOR [23], BoostThreader [22], SPARKS-X [24], MRFalign, and CEthreader [28]; (ii) machine learning methods such as FOLDpro [29], RF-Fold [31], DN-Fold [31], RFDN-Fold [31]; and (iii) deep learning methods such as DeepFR [38], CNN-BGRU [44] VGGfold [42], FoldTR [43], and FoldHSphere [45]. Table 4 shows the PFR accuracy results achieved by these methods on the LINDAHL test set, as well as the best performing model ProtT5 + LAT and the average ensemble. We notice that ProtT5 + LAT alone outperforms all state-of-the-art methods at the superfamily and fold levels. At the family level, it also obtains better accuracy than the rest of deep learning-based methods which, at this level, tend to perform worse than alignment methods. Furthermore, as we discussed in the previous section, the ensemble strategy shows a performance boost at the fold level, but also at the family level. At this level, it achieves a top 1 accuracy similar to the best alignment methods, and clearly outperforms all in top 5. Therefore, the use of the protein-LM embeddings as the input representation not only bridges the gap at the family and fold levels, but also increases the performance of fold recognition consistently at all levels.

**Table 4:**
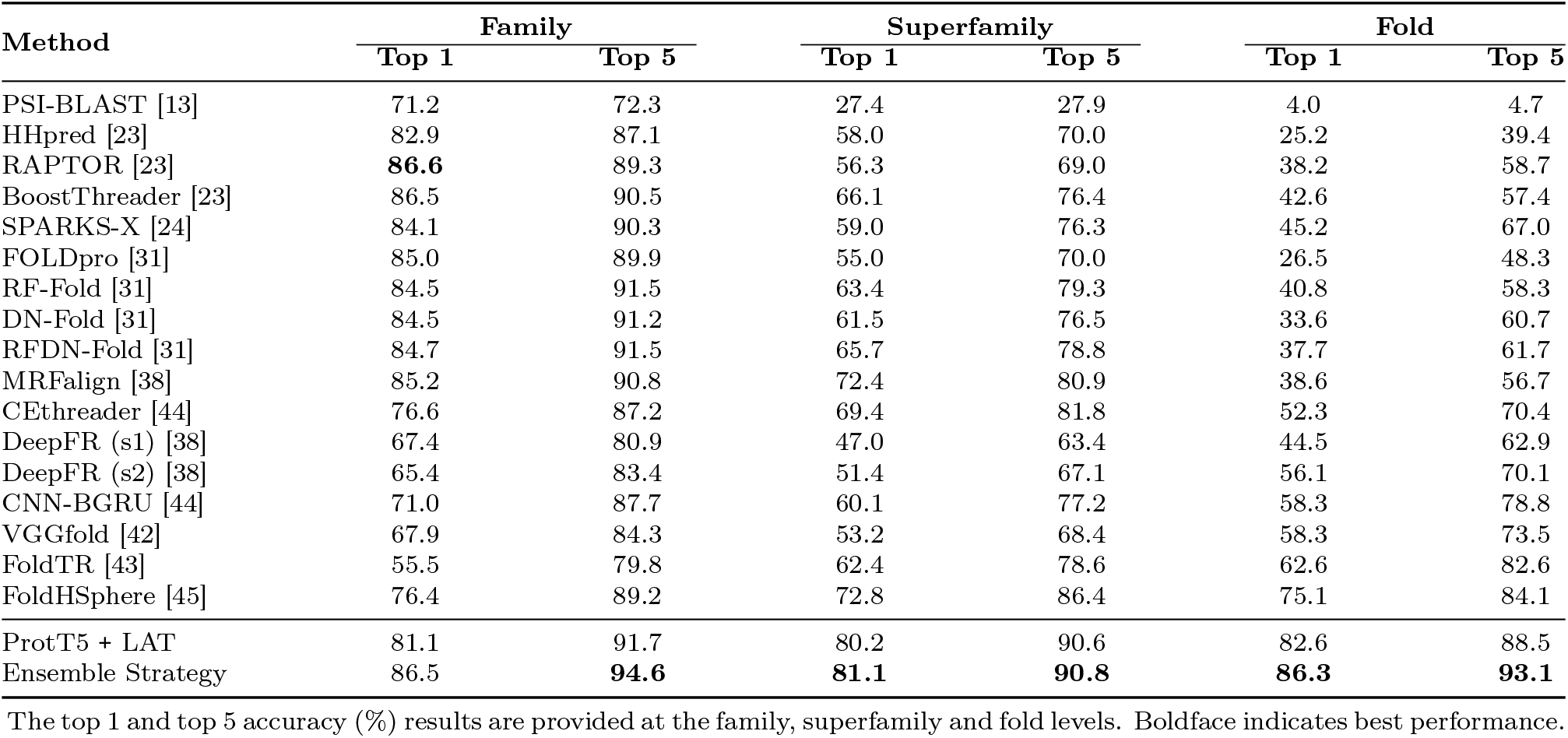
Three-level LINDAHL pairwise fold recognition (PFR) results of the best individual model and the ensembling strategy (average ensemble), in comparison with the state-of-the-art.

In addition, in order to compare with DeepFRpro [38], CNN-BGRU-RF+ [44], FoldTRpro [43], FSD_XGBoostpro [43], and FoldHSphere [45], which apply a random forest (RF) ensemble, we implemented and tested the same RF strategy in our ensemble approach (see Materials and Methods section). It must be noted that these results cannot be directly compared to the previous ones, as this approach involves additional training of the RF models on the test set in a 10-stage cross-testing manner. From Table 5 we can see that the RF ensemble introduced here consistently outperforms all previous state-of-the-art methods at the superfamily and fold levels. However, it provides lower accuracy at the family level. Interestingly, this evaluation scenario seems to lead to unexpected results due to cross-testing. That is why we believe average ensembling provides more reliable results than the RF ensemble, and can be compared more fairly against the individual models.

**Table 5:** Three-level LINDAHL pairwise fold recognition (PFR) results of the random forest (RF) ensemble, in comparison with the state-of-the-art.

| Method | Family |  | Superfamily |  | Fold |  |
| --- | --- | --- | --- | --- | --- | --- |
|  | Top 1 | Top 5 | Top 1 | Top 5 | Top 1 | Top 5 |
| DeepFRpro (s1) [38] | **85.6** | 91.9 | 66.6 | 82.0 | 57.6 | 73.8 |
| DeepFRpro (s2) [38] | 83.1 | 92.3 | 69.6 | 82.5 | 66.0 | 78.8 |
| CNN-BGRU-RF+ [44] | 85.4 | 93.5 | 73.3 | 87.8 | 76.3 | 85.7 |
| FoldTRpro [43] | 83.8 | 92.8 | 76.0 | 89.2 | 58.3 | 87.2 |
| FSD_XGBoostpro [43] | 82.7 | **94.6** | 77.9 | 91.5 | 68.2 | 91.9 |
| FoldHSpherePro [45] | 85.2 | 93.0 | 79.0 | 89.2 | 81.3 | 90.3 |
| RF Ensemble | 79.6 | 92.8 | **82.5** | **92.4** | **90.3** | **94.4** |

The top 1 and top 5 accuracy (%) results are provided at the family, superfamily and fold levels. Boldface indicates best performance.

For the DFC task we compare to deep learning methods that allow for the direct classification of protein domains into different folds. In Table 6 we show the results for the LINDAHL_1.75 and SCOP_2.06 test sets. Using both sets we evaluated the state-of-the-art DeepFR, CNN-BGRU, and FoldHSphere methods. In addition, for SCOP_2.06 we include the results of the DeepSF method [57], which originally introduced the SCOP_2.06 set and the DFC task. In the case of LINDAHL_1.75 (Table 6a), we observe that the top-performing ProtT5 + RBG model obtains better results than previous methods at all three levels, considerably outperforming them at the superfamily and fold levels. For SCOP_2.06 (Table 6b), the top-performing ESM-MSA + RBG model seems to fit the family and superfamily subsets almost perfectly, obtaining accuracy values above 99%. It also generalizes better than previous methods in the fold subset, with a top 1 accuracy of 70.3% (81.1% top 5) which is more than 15 percentage points higher than the previous methods DeepSF and FoldHSphere (both obtained 51.5% top 1 and 65.7% top 5 accuracies). Moreover, as discussed before, the ensemble strategy already outperformed the best individual models on both test sets and, therefore, all the considered methods from the state-of-the-art.

**Table 6:** Three-level **(a)** LINDAHL_1.75 and **(b)** SCOP_2.06 direct fold classification (DFC) results of the best individual models and the ensembling strategy (soft voting ensemble), in comparison with the state-of-the-art.

| (a) LINDAHL_1.75 Test Set |  |  |  |  |  |  |  |  |
| --- | --- | --- | --- | --- | --- | --- | --- | --- |
| Method | Full Set |  | Family |  | Superfamily |  | Fold |  |
|  | Top 1 | Top 5 | Top 1 | Top 5 | Top 1 | Top 5 | Top 1 | Top 5 |
| DeepFR (s1) | 57.3 | 72.5 | 70.7 | 83.9 | 34.8 | 55.7 | 11.4 | 25.7 |
| DeepFR (s2) | 66.5 | 76.0 | 79.5 | 86.5 | 45.2 | 57.6 | 20.0 | 42.9 |
| CNN-BGRU | 70.3 | 85.1 | 80.7 | 92.7 | 48.6 | 71.0 | 47.1 | 62.9 |
| FoldHSPHERE | 82.0 | 90.9 | 92.7 | 97.3 | 66.2 | 81.4 | 38.6 | 65.7 |
| ProtT5 + RBG | 87.6 | 92.4 | 94.3 | 97.1 | 79.5 | 88.1 | 55.7 | 65.7 |
| Ensemble Strategy | **92.3** | **97.5** | **97.6** | **99.3** | **89.1** | **97.6** | **57.1** | **81.4** |

| (b) SCOP_2.06 Test Set |  |  |  |  |  |  |  |  |
| --- | --- | --- | --- | --- | --- | --- | --- | --- |
| Method | Full Set |  | Family |  | Superfamily |  | Fold |  |
|  | Top 1 | Top 5 | Top 1 | Top 5 | Top 1 | Top 5 | Top 1 | Top 5 |
| DeepFR (s1) | 89.2 | 94.2 | 88.1 | 93.9 | 91.1 | 95.4 | 24.3 | 37.8 |
| DeepFR (s2) | 91.2 | 95.6 | 91.1 | 95.8 | 92.3 | 96.4 | 37.8 | 54.1 |
| DeepSF [57] | 73.0 | 90.3 | 75.9 | 91.8 | 72.2 | 90.1 | 51.4 | 67.6 |
| CNN-BGRU | 89.9 | 96.5 | 91.5 | 96.8 | 90.1 | 97.0 | 48.7 | 64.9 |
| FoldHSPHERE | 96.5 | 98.3 | 97.2 | 98.4 | 97.1 | 98.9 | 51.4 | 67.6 |
| ESM-MSA + RBG | 99.2 | 99.4 | **99.5** | 99.5 | 99.7 | 99.8 | 70.3 | 81.1 |
| Ensemble Strategy | **99.3** | **99.6** | 99.3 | **99.6** | **99.8** | **99.9** | **75.7** | **86.5** |

The top 1 and top 5 accuracy (%) results are provided for the full test set, and the family, superfamily and fold subsets. Boldface indicates best performance per set.

Taken together, these results show the superiority of protein-LM embeddings over other sequence representations such as the PSSM and secondary structure (DeepSF, CNN-BGRU, and FoldHSphere), or two-dimensional representations such as contact maps (DeepFR, VGGfold, and FoldTR).

## 4 Conclusion

This work provides a comparative analysis of different pre-trained embeddings from protein language models (protein-LMs) in protein fold prediction. To do so, we fine-tuned the protein-LM embeddings (*LMEmb*) through several state-of-the-art neural network models using fold labels for supervised training. The performance was evaluated in two differentiated tasks: pairwise fold recognition (PFR) and direct fold classification (DFC). For the PFR task, we extracted a fold-representative embedding vector (*FoldEmb*), which we used to predict the protein fold class by pairwise comparison with known protein domains. In contrast, in the DFC task we directly mapped the protein domain into one of the more than thousand existing fold classes in the SCOP database. For both tasks, the protein-LM embeddings learned by pre-trained transformer-based models proved to be more effective at identifying the protein fold than those extracted from LSTM-based models. Thus, the ESM-MSA and ProtT5 amino acid-level embeddings in combination with the RBG and LAT architectures provided the best PFR and DFC results. Compared to the state-of-the-art, these models alone were able to predict the fold class with higher accuracy at the three levels—family, superfamily and fold. Furthermore, our proposed ensemble approaches provided a significant performance boost over these individual models and thus over the current state-of-the-art. These results demonstrate the suitability of protein-LM embeddings over other traditional protein representations for fold prediction.

## Footnotes

1 https://github.com/songlab-cal/tape

2 https://github.com/mheinzinger/SeqVec

3 https://github.com/facebookresearch/esm

4 https://github.com/agemagician/ProtTrans

