## Supplementary Material for "An Analysis of Protein Language Model Embeddings for Fold Prediction"

Amelia Villegas-Morcillo, Angel M. Gomez, Victoria Sanchez

Department of Signal Theory, Telematics and Communications,
University of Granada, Spain

**Table S1:** Performance of the *LMEmb* embeddings in the pairwise fold recognition (PFR) task, using the LINDAHL test set. The top 1 and top 5 accuracy (%) results are provided at the family, superfamily and fold levels. We compare the performance of the amino acid-level embeddings *LMEmb-AA* and protein-level embeddings *LMEmb-Prot*, using $L_1$ distance as comparison metric. Underline indicates best performance.

| Embeddings | Family | | Superfamily | | Fold | |
|---|---|---|---|---|---|---|
| | Top 1 | Top 5 | Top 1 | Top 5 | Top 1 | Top 5 |
| ***LMEmb-AA*** ($L \times F$) | | | | | | |
| UniRep | 51.4 | 64.0 | 30.0 | 41.7 | 17.1 | 32.1 |
| SeqVec | 31.7 | 45.4 | 5.8 | 12.9 | 0.3 | 4.4 |
| ESM-1b | 60.2 | 66.5 | 14.5 | 22.4 | 3.7 | 9.0 |
| ESM-MSA | <u>83.1</u> | <u>90.1</u> | <u>55.8</u> | 64.5 | 20.2 | 35.8 |
| ProtBERT | 33.2 | 44.3 | 6.0 | 12.0 | 3.4 | 13.4 |
| ProtT5 | 74.6 | 84.9 | 25.6 | 40.6 | 4.7 | 13.7 |
| ***LMEmb-Prot*** ($F$) | | | | | | |
| UniRep | 45.6 | 62.5 | 34.1 | 45.6 | 19.6 | 36.8 |
| SeqVec | 59.1 | 75.0 | 39.9 | 53.9 | 16.8 | 36.1 |
| ESM-1b | 82.2 | 89.0 | 47.0 | <u>65.7</u> | <u>21.2</u> | <u>39.6</u> |
| ESM-MSA | 76.9 | 88.3 | 43.3 | 54.4 | 15.9 | 29.3 |
| ProtBERT | 45.9 | 62.0 | 13.8 | 28.8 | 9.7 | 20.6 |
| ProtT5 | 79.8 | 89.4 | 35.0 | 56.0 | 16.2 | 30.5 |

**Table S2:** Performance of the *LMEmb-Prot* embeddings in the direct fold classification (DFC) task, using the **(a)** LINDAHL_1.75 and **(b)** SCOP_2.06 test sets, and cosine similarity as comparison metric. The top 1 and top 5 accuracy (%) results are provided for the full test set, and the family, superfamily and fold subsets. Underline indicates best performance per set.

**(a)** LINDAHL_1.75 Test Set

| Embeddings | Full Set | | Family | | Superfamily | | Fold | |
|---|---|---|---|---|---|---|---|---|
| | Top 1 | Top 5 | Top 1 | Top 5 | Top 1 | Top 5 | Top 1 | Top 5 |
| UniRep | 42.1 | 53.2 | 49.4 | 61.8 | 29.1 | 35.7 | 20.0 | 32.9 |
| SeqVec | 57.1 | 66.7 | 66.8 | 77.0 | 42.4 | 50.5 | 18.6 | 28.6 |
| ESM-1b | <u>73.8</u> | <u>80.0</u> | 83.1 | <u>88.5</u> | <u>62.9</u> | <u>69.1</u> | <u>28.6</u> | <u>41.4</u> |
| ESM-MSA | 68.4 | 73.7 | <u>83.3</u> | 87.5 | 43.8 | 53.3 | 17.1 | 18.6 |
| ProtBERT | 28.7 | 43.6 | 34.7 | 52.3 | 17.6 | 27.1 | 11.4 | 20.0 |
| ProtT5 | 68.7 | 76.7 | 79.7 | 87.5 | 51.9 | 61.0 | 25.7 | 32.9 |

**(b)** SCOP_2.06 Test Set

| Embeddings | Full Set | | Family | | Superfamily | | Fold | |
|---|---|---|---|---|---|---|---|---|
| | Top 1 | Top 5 | Top 1 | Top 5 | Top 1 | Top 5 | Top 1 | Top 5 |
| UniRep | 59.1 | 69.6 | 78.6 | 84.8 | 51.9 | 63.7 | 10.8 | 43.2 |
| SeqVec | 77.9 | 84.6 | 91.8 | 93.9 | 73.2 | 81.4 | 21.6 | 54.1 |
| ESM-1b | 94.0 | 96.6 | 95.7 | 97.7 | 94.6 | 97.0 | 35.1 | 54.1 |
| ESM-MSA | <u>95.9</u> | <u>96.8</u> | <u>96.9</u> | 97.6 | <u>96.9</u> | <u>97.6</u> | 27.0 | 43.2 |
| ProtBERT | 57.5 | 70.3 | 83.2 | 89.1 | 47.4 | 62.8 | 21.6 | 48.7 |
| ProtT5 | 93.9 | <u>96.8</u> | 95.4 | <u>98.0</u> | 94.0 | 96.8 | <u>56.8</u> | <u>75.7</u> |

**Table S3:** Pairwise fold recognition (PFR) top 1 accuracy (%) results on the LINDAHL test set. At each level (family, superfamily and fold), we compare the performance of the 6 protein-LM embeddings and the 3 neural architectures trained using either softmax cross-entropy loss or large margin cosine loss (LMCL). Underline indicates best performance for each loss function and model, boldface indicates best overall.

| Embeddings | MLP Model | | | RBG Model | | | LAT Model | | |
|---|---|---|---|---|---|---|---|---|---|
| | Family | Superfamily | Fold | Family | Superfamily | Fold | Family | Superfamily | Fold |
| **Softmax Loss** | | | | | | | | | |
| UniRep | 49.5 | 38.9 | 28.0 | 36.6 | 40.1 | 37.4 | 34.1 | 37.6 | 29.0 |
| SeqVec | 64.3 | 50.7 | 34.6 | 45.8 | 46.3 | 43.0 | 61.4 | 48.6 | 42.4 |
| ESM-1b | 82.3 | 71.4 | 58.9 | 64.1 | 62.4 | 60.7 | 81.6 | 77.2 | 63.9 |
| ESM-MSA | 85.6 | 63.8 | 34.6 | 82.0 | 76.3 | 67.9 | 82.3 | 79.7 | 58.3 |
| ProtBERT | 68.5 | 45.2 | 29.6 | 56.4 | 54.8 | 49.5 | 69.7 | 54.8 | 30.8 |
| ProtT5 | **86.7** | 77.0 | 60.1 | 68.5 | 68.2 | 73.5 | **85.0** | **81.1** | 64.5 |
| **LMCL** | | | | | | | | | |
| UniRep | 48.5 | 42.6 | 41.1 | 46.5 | 45.4 | 45.8 | 45.0 | 40.1 | 40.2 |
| SeqVec | 61.8 | 55.8 | 51.1 | 53.3 | 52.1 | 48.9 | 59.1 | 59.2 | 60.1 |
| ESM-1b | 80.9 | 74.7 | **73.5** | 75.1 | 74.2 | 74.1 | 82.0 | 76.3 | 74.5 |
| ESM-MSA | 85.2 | 72.1 | 60.4 | **83.2** | **81.3** | 73.8 | 78.0 | 75.8 | 75.4 |
| ProtBERT | 65.0 | 57.8 | 49.5 | 67.7 | 69.4 | 66.7 | 67.0 | 63.8 | 61.1 |
| ProtT5 | 82.9 | **78.6** | **73.5** | 79.3 | 79.7 | **80.4** | 81.1 | 80.2 | **82.6** |

**Table S4:** Ensemble results for the pairwise fold recognition (PFR) task using LINDAHL. Here we include the results of the 3 best performing protein-LM embeddings (ESM-1b, ESM-MSA, and ProtT5) with 2 neural architectures (RBG and LAT); as well as the average ensemble for different combinations of the 6 individual models. For each one, the top 1 and top 5 accuracy (%) results are provided at the family, superfamily and fold levels. Underline indicates best performance per group, boldface indicates best overall.

| Embeddings | Models | Family | | Superfamily | | Fold | |
|---|---|---|---|---|---|---|---|
| | | Top 1 | Top 5 | Top 1 | Top 5 | Top 1 | Top 5 |
| **Individual Models** | | | | | | | |
| ESM-1b | RBG | 75.1 | 87.2 | 74.2 | 84.6 | 74.1 | 83.2 |
| | LAT | 82.0 | 90.6 | 76.3 | 86.4 | 74.5 | 84.1 |
| ESM-MSA | RBG | 83.2 | 93.2 | 81.3 | 87.3 | 73.8 | 87.2 |
| | LAT | 78.0 | 91.9 | 75.8 | 86.2 | 75.4 | 85.0 |
| ProtT5 | RBG | 79.3 | 91.7 | 79.7 | 88.5 | 80.4 | 88.5 |
| | LAT | 81.1 | 91.7 | 80.2 | 90.6 | 82.6 | 88.5 |
| **Average Ensemble** | | | | | | | |
| ESM-1b | RBG + LAT | 81.4 | 89.9 | 75.3 | 84.8 | 75.4 | 86.0 |
| ESM-MSA | RBG + LAT | 84.1 | 93.9 | 80.2 | 86.6 | 76.9 | 88.5 |
| ProtT5 | RBG + LAT | 81.4 | 92.4 | 80.2 | 91.0 | 84.7 | 89.7 |
| ESM-1b + ESM-MSA + ProtT5 | RBG | 82.9 | 93.5 | **83.6** | **91.2** | 84.7 | 92.5 |
| | LAT | **87.2** | **95.0** | 80.9 | 90.8 | 82.9 | 91.0 |
| | RBG + LAT | 86.5 | 94.6 | 81.1 | 90.8 | **86.3** | **93.1** |

**Table S5:** Ensemble results for the direct fold classification (DFC) task using **(a)** LINDAHL_1.75 and **(b)** SCOP_2.06. Here we include the results of the 3 best performing protein-LM embeddings (ESM-1b, ESM-MSA, and ProtT5) with 2 neural architectures (RBG and LAT); as well as the soft voting ensemble for different combinations of the 6 individual models. For each one, the top 1 and top 5 accuracy (%) results are provided for the full test set, and the family, superfamily and fold subsets. Underline indicates best performance per group, boldface indicates best overall.

**(a)** LINDAHL_1.75 Test Set

| Embeddings | Model | Full Set | | Family | | Superfamily | | Fold | |
|---|---|---|---|---|---|---|---|---|---|
| | | Top 1 | Top 5 | Top 1 | Top 5 | Top 1 | Top 5 | Top 1 | Top 5 |
| **Individual Models** | | | | | | | | | |
| ESM-1b | RBG | 83.7 | 89.0 | 92.2 | 95.3 | 74.3 | 82.9 | 40.0 | 54.3 |
| | LAT | 85.7 | 89.7 | 92.6 | 94.6 | 81.9 | 86.7 | 38.6 | 57.1 |
| ESM-MSA | RBG | 86.6 | 93.3 | 96.5 | 98.1 | 74.3 | 89.1 | 40.0 | 65.7 |
| | LAT | 84.6 | 91.4 | 94.4 | 97.6 | 71.4 | 83.8 | 41.4 | 61.4 |
| ProtT5 | RBG | 87.6 | 92.4 | 94.3 | 97.1 | 79.5 | 88.1 | 55.7 | 65.7 |
| | LAT | 87.6 | 94.3 | 95.6 | 97.8 | 77.6 | 91.9 | 50.0 | 71.4 |
| **Soft Voting Ensemble** | | | | | | | | | |
| ESM-1b | RBG + LAT | 85.9 | 90.7 | 93.7 | 96.1 | 77.6 | 86.2 | 44.3 | 58.6 |
| ESM-MSA | RBG + LAT | 88.1 | 94.5 | 97.0 | 97.8 | 79.1 | 91.9 | 40.0 | 74.3 |
| ProtT5 | RBG + LAT | 89.8 | 94.4 | 96.3 | 98.0 | 81.9 | 91.9 | **58.6** | 71.4 |
| ESM-1b + ESM-MSA + ProtT5 | RBG | 92.2 | 97.0 | **97.8** | **99.3** | 88.1 | 96.7 | 57.1 | 78.6 |
| | LAT | 90.8 | 96.3 | 97.5 | **99.3** | 85.7 | 95.7 | 50.0 | 72.9 |
| | RBG + LAT | **92.3** | **97.5** | 97.6 | **99.3** | **89.1** | **97.6** | 57.1 | **81.4** |

**(b)** SCOP_2.06 Test Set

| Embeddings | Model | Full Set | | Family | | Superfamily | | Fold | |
|---|---|---|---|---|---|---|---|---|---|
| | | Top 1 | Top 5 | Top 1 | Top 5 | Top 1 | Top 5 | Top 1 | Top 5 |
| **Individual Models** | | | | | | | | | |
| ESM-1b | RBG | 96.1 | 97.6 | 97.6 | 98.4 | 96.5 | 98.1 | 48.7 | 62.2 |
| | LAT | 96.5 | 97.2 | 97.8 | 98.4 | 96.9 | 97.6 | 51.4 | 56.8 |
| ESM-MSA | RBG | 99.2 | 99.4 | **99.5** | 99.5 | 99.7 | 99.8 | 70.3 | 81.1 |
| | LAT | 98.9 | 99.4 | 99.1 | 99.3 | 99.4 | 99.8 | 70.3 | 83.8 |
| ProtT5 | RBG | 97.8 | 98.9 | 98.3 | 98.8 | 98.4 | 99.4 | 59.5 | 75.7 |
| | LAT | 98.0 | 99.3 | 98.3 | 99.2 | 98.5 | 99.7 | 67.6 | 83.8 |
| **Soft Voting Ensemble** | | | | | | | | | |
| ESM-1b | RBG + LAT | 97.0 | 98.2 | 98.4 | 99.1 | 97.4 | 98.5 | 51.4 | 67.6 |
| ESM-MSA | RBG + LAT | 99.1 | 99.5 | 99.3 | 99.5 | 99.5 | **99.9** | **75.7** | 83.8 |
| ProtT5 | RBG + LAT | 98.5 | 99.2 | 98.9 | 99.3 | 98.8 | 99.5 | 73.0 | 83.8 |
| ESM-1b + ESM-MSA + ProtT5 | RBG | **99.3** | **99.6** | 99.3 | **99.6** | **99.8** | **99.9** | 73.0 | 83.8 |
| | LAT | 99.2 | **99.6** | 99.3 | **99.6** | 99.7 | **99.9** | 70.3 | 83.8 |
| | RBG + LAT | **99.3** | **99.6** | 99.3 | **99.6** | **99.8** | **99.9** | **75.7** | **86.5** |

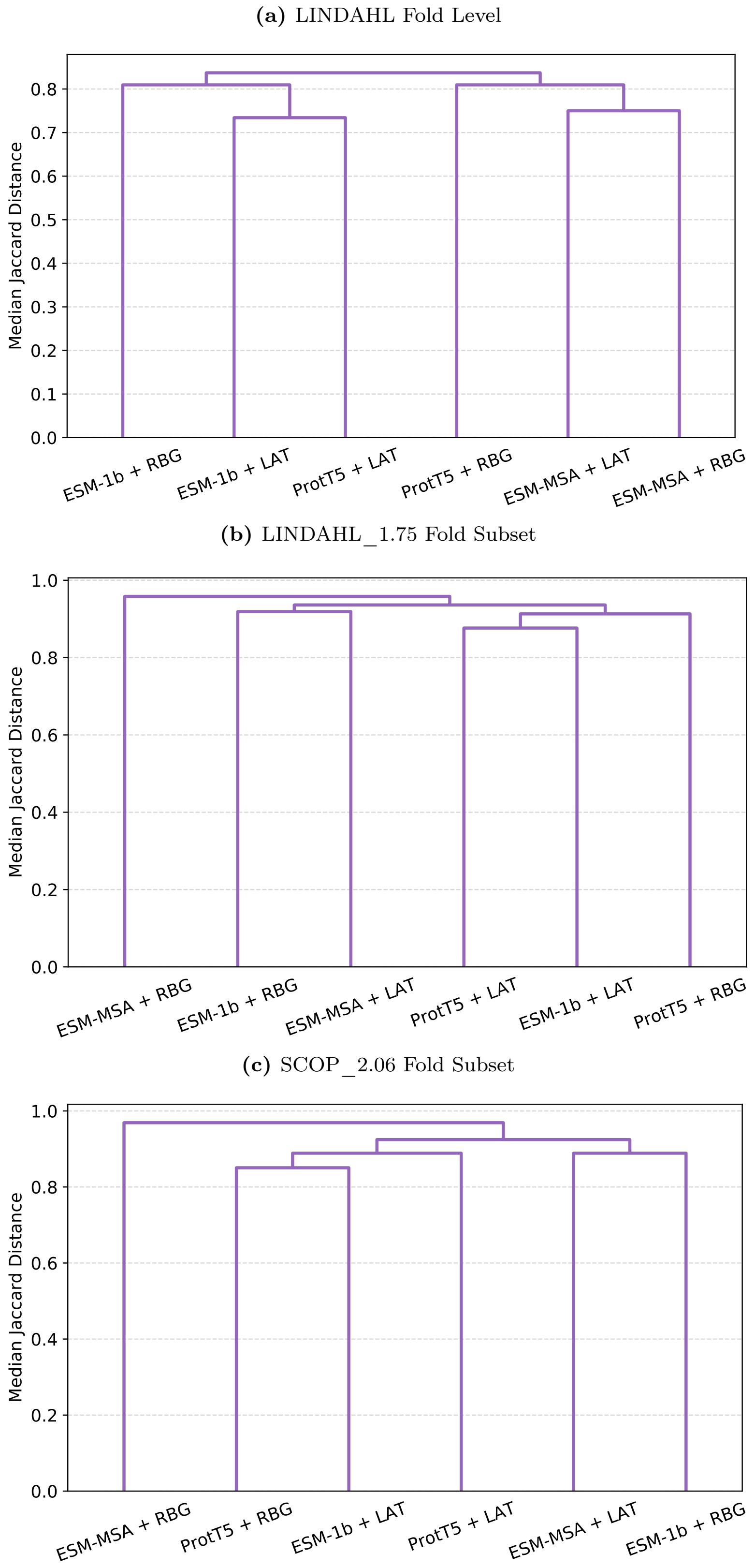


**Figure S1:** Hierarchical clustering with complete linkage of the 6 best performing models (ESM-1b, ESM-MSA, and ProtT5 embeddings with RBG and LAT architectures). As metric, we used the median of the Jaccard distance distribution between each two models. **(a)** For the LINDAHL test set (PFR task), we compared the 50 closest neighbors of each protein domain at the fold level, using the cosine similarity scores of their fold-representative embeddings. For the **(b)** LINDAHL_1.75 and **(c)** SCOP_2.06 test sets (DFC task), we compared the 50 fold classes with maximum score for each domain in the fold subset. In all cases, the models cluster at very high values of median Jaccard distance, suggesting that the scores provided by each model are quite dissimilar.
